# Genome-wide interaction study of a proxy for stress-sensitivity and its prediction of major depressive disorder

**DOI:** 10.1101/194290

**Authors:** Aleix Arnau-Soler, Mark J. Adams, Generation Scotland, Major Depressive Disorder Working Group of the Psychiatric Genomics Consortium, Caroline Hayward, Pippa A. Thomson

## Abstract

Individual response to stress is correlated with neuroticism and is an important predictor of both neuroticism and the onset of major depressive disorder (MDD). Identification of the genetics underpinning individual differences in response to negative events (stress-sensitivity) may improve our understanding of the molecular pathways involved, and its association with stress-related illnesses. We sought to generate a proxy for stress-sensitivity through modelling the interaction between SNP allele and MDD status on neuroticism score in order to identify genetic variants that contribute to the higher neuroticism seen in individuals with a lifetime diagnosis of depression compared to unaffected individuals. Meta-analysis of genome-wide interaction studies (GWIS) in UK Biobank (N = 23,092) and Generation Scotland: Scottish Family Health Study (N = 7,155) identified no genome-wide significance SNP interactions. However, gene-based tests identified a genome-wide significant gene, *ZNF366*, a negative regulator of glucocorticoid receptor function implicated in alcohol dependence (*p* = 1.48×10^-7^; Bonferroni-corrected significance threshold *p* < 2.79×10^-6^). Using summary statistics from the stress-sensitivity term of the GWIS, SNP heritability for stress-sensitivity was estimated at 5.0%. In models fitting polygenic risk scores of both MDD and neuroticism derived from independent GWAS, we show that polygenic risk scores derived from the UK Biobank stress-sensitivity GWIS significantly improved the prediction of MDD in Generation Scotland. This study may improve interpretation of larger genome-wide association studies of MDD and other stress-related illnesses, and the understanding of the etiological mechanisms underpinning stress-sensitivity.

## Introduction

Stressful life events are known to increase liability to mental illness and disease-related traits [1] including neuroticism [2-4], major depressive disorder (MDD) [5-7], autoimmune diseases [8] and some cancers [9, 10]. A greater understanding of the causal mechanism by which negative events affect disease risk or outcome may be beneficial in identifying individuals for targeted support. However, it has been proposed that sensitivity to stress may be an important predictor of response to stress [11, 12]. In particular, the effect on an individual may result more from the perceived stress than the event itself, and may be dependent on individual differences in stress-sensitivity [13-18]. Studies of 5-HTT and twin studies suggest that stress-sensitivity may, at least in part, be heritable [19-22]. Despite a complex interaction between MDD, neuroticism and stress, multivariate structural equation models have confirmed a genetic effect on perceived stress, overlapping that on MDD or neuroticism, but with a specific genetic component [21]. The inter-relatedness of these traits may offer an approach to identify the genetic variation that affects an individual’s stress-sensitivity, and improve genetic prediction of an individual’s liability to negative outcomes. By modelling the interaction between SNP allele and MDD status on neuroticism score through genome-wide interaction studies (GWIS), we sought to investigate the genetics of stress-sensitivity.

The personality trait neuroticism is moderately heritable (30–50% estimates from twin studies) [23-26], is higher in individuals with depression compared to controls [27, 28] and is known to have shared genetic aetiology with depression [29-32]. Neuroticism is strongly correlated with measures of sensitivity to punishment but not reward [33], positively correlated with perceived personal relevance of a stressor [34, 35] and has been used previously as a proxy measure of stress-sensitivity [36]. Neuroticism is thought to mediate or interact with the effects of adverse life events on risk of depression [5, 37]. It has a substantial stable component [38], however, there is evidence for change, as well as stability, across the life span [2-4, 39]. Individual differences in neuroticism are enduringly influenced by both genetic and environmental factors [40]. Whereas the stable component of neuroticism is strongly determined by genetics, change in neuroticism score is attributed to the effects of unshared environment [39]. Persistent change in neuroticism score has been shown in response to life events [2-4]. Negative life events lead to small persistent increases in neuroticism over time [3]. However, recent stressful life events (β = 0.14 95%CI 0.13 - 0.15, *p* < 0.001) have a stronger effect than distant stressful life events suggesting a reduction of effect over time [3]. Long-lasting increases in neuroticism associated with distant negative life events are mediated by depression [4].

Major depressive disorder (MDD) is a complex disorder influenced by both genetic contributions and environmental risk factors, with heritability estimates from twin and family studies of between 31-42% [41, 42]. Confirmed environmental risk factors for MDD include maternal infections, childhood maltreatment and negative life events [5-7, 43, 44]. However, few genetic studies have such information and even fewer prospective studies exist. Incorporation of stressful life events has been shown to improve the ability to predict MDD [45, 46] and, although stress is an environmental risk factor, it may have an independent genetic contribution to risk of depression [46-50].

These studies suggest that a genetic variable derived from the difference in neuroticism levels seen in individuals with MDD compared to controls may allow us to identify genetic loci important for stress-sensitivity. We sought to identify the genetic underpinnings of individual’s sensitivity to stress response (stress-sensitivity) by identifying variants that contribute to the higher neuroticism levels seen in individuals with a lifetime diagnosis of MDD. Further, polygenic risk scores (PRS) derived from this stress-sensitivity variable may improve prediction of MDD over that based on MDD or neuroticism PRS alone.

Using unrelated individuals from two large population-based samples, UK Biobank (UKB; N = 23,092) and Generation Scotland: Scottish Family Health Study (GS:SFHS; N = 7,155), we sought to identify genes involved in stress-sensitivity by performing GWIS for the interaction between MDD status and SNP allele on neuroticism score. We identified a gene significantly associated with stress-sensitivity and show that a PRS derived from the interaction term of the GWIS, significantly predicts liability to depression independently of the PRS for MDD and/or neuroticism.

## Materials and methods

### UK Biobank (UKB) Participants

UKB is a major national health resource that aims to improve the prevention, diagnosis and treatment of a wide range of illnesses. It recruited more than 500,000 participants aged from middle to older age who visited 22 assessment centres across the UK between 2006 and 2010. Data were collected on background and lifestyle, cognitive and physical assessments, sociodemographic factors and medical history. The scientific rationale, study design, ethical approval, survey methods, and limitations are reported elsewhere [51, 52]. UKB received ethical approval from the NHS National Research Ethics Service North West (Research Ethics Committee Reference Number: 11/NW/0382). All participants provided informed consent. The present study was conducted on genome-wide genotyping data available from the initial release of UKB data (released 2015). Details of sample processing specific to UKB project are available at http://biobank.ctsu.ox.ac.uk/crystal/refer.cgi?id=155583 and the Axiom array at http://media.affvmetrix.com/support/downloads/manuals/axiom_2_assay_auto_workflow_user_guide.pdf. UKB genotyping and the stringent QC protocol applied to UKB data before it was released can be found at http://biobank.ctsu.ox.ac.uk/crvstal/refer.cgi?id=155580. SNPs genotyped on GS:SFHS were extracted from the imputed UKB genotype data [53] (imputed by UKB using a merged panel of the UK10K haplotype reference panel and the 1000 Genomes Phase 3 reference panel) with quality > 0.9 was hard-called using PLINK v1.9 [54]. Individuals were removed based on UKB genomic analysis exclusion (UKB Data Dictionary item #22010), non-white British ancestry (#22006: genetic ethnic grouping; from those individuals who self-identified as British, principal component analysis was used to remove outliers), high genotype missingness (#22005), genetic relatedness (#22012; no pair of individuals have a KING-estimated kinship coefficient > 0.0442), QC failure in UK BiLEVE study (#22050 and #22051: UK BiLEVE Affymetrix and UK BiLEVE genotype quality controls for samples) and gender mismatch (#22001: genetic sex). Further, from the initial release of UKB data and using PLINK pi-hat < 0.05, individuals who were also participants of GS:SFHS and their relatives were excluded to remove any overlap of individuals between discovery and target samples. A dataset of 109,283 individuals with 557,813 SNPs remained for further analysis, aged 40-79 (57,328 female, 51,954 male; mean age = 57.1 years, s.d. = 7.99), of which 109,282 had data available for neuroticism score and 23,092 had data available on MDD status (*n*_cases_ = 7,834, *n*_controls_ = 15,258, *n*_female_ = 11,510, *n*_male_ = 11,582; mean age = 57.7 years, s.d. = 8.04). Thus, the final dataset comprised 23,092 unrelated individuals.

### Generation Scotland Scottish Family Health Study (GS:SFHS) Participants

GS:SFHS is a family-based genetic epidemiology study which includes 23,960 participants from ~ 7,000 Scottish family groups collected by a cross-disciplinary collaboration of Scottish medical schools and the National Health Service (NHS) from Feb 2006 to Mar 2011. Participants were interviewed and clinically assessed for a wide range of health-related traits (including high-fidelity phenotyping for Major Depressive Disorder and related endophenotypes), environmental covariates and linked to routine health records [55, 56]. All components of GS:SFHS obtained ethical approval from the Tayside Committee on Medical Research Ethics on behalf of the NHS (Research Ethics Committee Reference Number: 05/S1401/89) and participants provided written consent. The protocol for recruitment is described in detail in previous publications [57, 58]. GS:SFHS genotyping and quality control is detailed elsewhere [59]. Briefly, individuals with more than 2% missing genotypes and sex discrepancies were removed, as well as population outliers. SNPs with genotype missingness > 2%, minor allele frequency < 1% and a Hardy-Weinberg Equilibrium test *p* < 1×10^−6^ were exclude. Finally, individuals were removed based on relatedness (pi-hat < 0.05), maximizing retention of case individuals, using PLINK v1.9 [54]. Genome-wide SNP data for further analysis comprised 7,233 unrelated individuals genotyped for 560,698 SNPs (*n*_female_ = 3,476, *n*_male_ = 3,757; PLINK v1.9 [54]), aged 18-92 (mean age = 50.4 years, s.d. = 12.06) of which: 7,190 had clinical data on MDD; 7,196 individuals had data on neuroticism; and 7,155 had data on both neuroticism and MDD.

### Phenotype assessment

#### Neuroticism score (EPQN)

Participants in both UKB and GS:SFHS cohorts were assessed for neuroticism using 12 questions from the Eysenck Personality Questionnaire-Revised Short Form’s Neuroticism Scale (EPQN) [60-63]. Neuroticism can be scored by adding up the number of “Yes” responses on EPQN. This short scale has a reliability of more than 0.8 [64]. EPQN distributions were found to be sufficiently “normal” after assessment for skewness and kurtosis to be analysed using linear regression (both coefficients were between −1 and 1).

#### MDD diagnoses

In UKB, the MDD phenotype was derived following the definitions from Smith et al. [63] Current and previous depressive symptoms were assessed by items relating to the lifetime experience of minor and major depression [60], items from the Patient Health Questionnaire [65] and items on help-seeking for mental health [63]. Using a touchscreen questionnaire, participants were defined as probable cases if they i) answered “Yes” to the question “Ever depressed for a whole week” (UKB field: 4598), plus at least 2 weeks duration (UKB field: 4609), or ii) did report having seen a GP or psychiatrist for nerves, anxiety, tension or depression (UKB fields: 2090 and 2010) and reported symptoms (UKB field: 4631) with at least 2 weeks duration (UKB field: 5375). In our unrelated sample, 7,834 participants were diagnosed with MDD (with single, moderate or recurrent episodes) and 15,258 were controls (N = 23,092).

In GS:SFHS, participants took in-person clinical visits where they were screened for a history of psychiatric and emotional disorders (i.e., psychiatric, mood state/psychological distress, personality and cognitive assessment) by trained researchers using the Structured Clinical Interview for DSM-IV Non-Patient Version (SCID) [66], which is internationally validated to identify episodes of depression. Those participants that were positive in the initial screening continue through clinical interview and were administered the mood sections of the SCID. The SCID elicited the presence or absence of a lifetime history of MDD, age of onset and number of episodes. Participants fulfilling the criteria for at least one major depressive episode within the last month were defined as current MDD cases. Participants who were screened positive for Bipolar I Disorder were excluded. Those participants who were negative during the initial screening or did not fulfilled criteria for MDD were assigned as controls. Further details regarding the diagnostic assessment are reported elsewhere [56, 57]. All interviewers were trained for the administration of the SCID. Inter-rater reliability for the presence or absence of a lifetime diagnosis of major depressive disorder was good (Kappa = 0.86, *p* < 0.001, 95%CI 0.7 to 1.0). In our unrelated GWIS sample (N = 7,155), 2,010 had a lifetime diagnosis of MDD and 5,145 were controls.

### Statistical Methods

#### GWIS and derivation of a genetic stress-sensitivity effect

The effect size of an stress-sensitivity effect (*β_SS_*) was derived by performing a GWIS for the effect of the MDD status and SNP allele on EPQN (dependent variable) in both UKB and GS:SFHS cohorts using PLINK 1.90 (PLINK-command --gxe; fitting MDD diagnosis as a binary “group” effect) [54]. PLINK-command --gxe estimates the difference in allelic association with a quantitative trait (EPQN) between two groups (MDD cases vs. controls) producing effect estimates on each group and a test of significance for the interaction between SNP allele and MDD status. The interaction *p* value reflects the difference between the regression coefficient of the allelic effect in a linear model for EPQN in MDD cases (*β_A_*) and the same regression coefficient in a linear model for EPQN in controls (*β_B_*). The stress-sensitivity interaction effect was defined as the difference in allele effect between MDD cases and control groups.

Considering one SNP, the effect it confers to EPQN can be modelled by MDD status (control = 0, MDD case = 1) as follows:

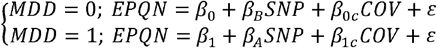

This is equivalent to modelling the effect on MDD cases as follows:

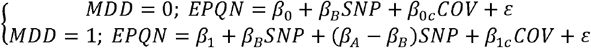

Or, it can be modelled as a whole as:

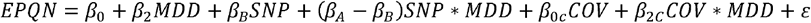

Where *COV* stands for covariates, *β*_2_ stands for *β*_1_ − *β*_0_, and *β*_2*c*_ stands for *β*_1*c*_ − *β*_0*c*_.

Thus, the interaction effect (*β_SS_*) can be estimated as the difference in allelic effect on EPQN between MDD cases (*β_A_*) and controls (*β_B_*) as follows,

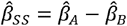

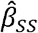 is therefore defined as the effect size reflecting the genetic stress-sensitivity effect on MDD cases compared to controls (S1 Fig).

#### Stress-sensitivity GWIS, main additive effect GWASs, meta-analysis and gene-set analysis

For GWIS and subsequent analyses, sample specific covariates were applied as follows: *UKB*. All phenotypes were adjusted for centre, array and batch as random effects prior to analyses. Analyses were adjusted for age, sex and 15 informative principal components (PCs; UKB Data Dictionary items #22009.01 to #22009.15) as fixed effects to take account of possible population stratification. *GS:SFHS*. All the analyses were adjusted for age, sex and 20 PCs.

GWAS for MDD and neuroticism, using logistic and linear models of additive allelic effects respectively, were conducted on the same sample sets for comparison and generation of matched PRS using PRSice-2 [67].

Results from the GWIS of UKB and GS:SFHS were combined in a sample size weighted meta-analysis performed using METAL [68]. While the use of standard error weighting is more common, the different diagnostic scheme and MDD prevalence between the two cohorts (GS:SFHS; 12.2%, UKB: 25.8%) [57, 63] may indicate systematic differences in the measurement of MDD. Generalized gene-based analysis of the meta-analysis was performed using MAGMA [69] implemented through FUMA[70] (http://fuma.ctglab.nl). Briefly, SNP summary statistics were mapped to 17,931 protein-coding genes. Individual SNP *p* values from a gene were combined into a gene test-statistic using a SNP-wise model and a known approximation of the sampling distribution used to obtain a gene-based *p* value. Genome-wide significance was defined at *p* = 0.05/17,931 = 2.79×10^-6^.

#### LD Score regression

The summary statistics from the meta-analysis were used to examine the genetic overlap between the polygenic architecture of stress-sensitivity, MDD and neuroticism. LD score regression was used to derive the genetic correlations (r_G_) between these traits [71, 72] using meta-analysed GWAS and GWIS summary statistics. SNP-based heritability was also estimated using LD score regression, using the summary statistics from single-SNP analyses.

#### Pathway, functional and gene expression analyses

Lead SNPs, independently associated with the phenotype, were identified using PLINK 1.90 by clumping (*p* threshold < 2×10^-5^; LD r2 > 0.1; physical kb threshold = 500kb; 1000 Genomes Project Phase 1 CEU, GBR, TSI genotype data), and analysed using DEPICT [73]. Further detail is given in ‘DEPICT analyses’ in S1 Supporting Information.

Genes associated with lead SNPs were investigated for evidence of: phenotypic association in the NCBI dbGaP database of genotypes and phenotypes [74] (https://www.ncbi.nlm.nih.gov/gap/phegeni), regulatory DNA elements in normal cell lines and association with expression quantitative trait loci (eQTLs) using the RegulomeDB database [75] (http://www.regulomedb.org) and the Genotype-Tissue Expression (GTEx) Portal [76] (http://www.gtexportal.org).

#### Polygenic profiling

PRS were produced using PRSice-2 [67], permuted 10,000 times and standardized to a mean of 0 and a standard deviation of 1. Using GWIS summary statistics, we created PRS for stress-sensitivity (PRS_SS_) by weighting the sum of the reference alleles in an individual by the stress-sensitivity effect (*β*_SS_). Additional PRS were generated weighting by MDD main additive effects (PRS_D_) and neuroticism main additive effects (PRS_N_) using GWAS summary statistics from GS:SFHS or UKB. In addition, PRS_D_ and PRS_N_ were also generated using summary statistics from the most recent Psychiatric Genetic Consortium (PGC) MDD meta-analysis [42] (excluding GS:SFHS, and UKB individuals when required; N = 155,866 & 138,884) and the Genetics of Personality Consortium (GPC) neuroticism meta-analysis [24, 77] (N = 63,661). Generalized linear models were implemented in R 3.1.3 [78]. The direct effect of PRS_SS_ (model 1), PRS_D_ (model 2) and PRS_N_ (model 3) on MDD risk were assessed in independent logistic regression models on GS:SFHS (target cohort) using GWAS and GWIS statistics from UKB (the largest cohort) as the discovery sample to weight PRS. Multiple regression models fitting both PRS_D_ and PRS_N_ (model 4) and fitting each of them separately with PRS_SS_ (models 5 and 6) were also calculated. Finally, full additive multiple regression models fitting PRS weighted by all three effects (full model) was assessed using both PRS_SS_, PRS_D_ and PRS_N_ at their best-fit in independent models. Further, results were also assessed using PRS_D_ and PRS_N_ weighted by PGC2 MDD [42] and GPC neuroticism [77] summary statistics. Further detail is given in ‘Polygenic Profiling’ in S1 Supporting Information. All models were adjusted by sex, age and 20 PCs. A null model was estimated from the direct effects of all covariates on MDD. 10,000 permutations were used to assess significance of each PRS. The predictive improvement of combining the effects of multiple PRS over a single PRS alone was tested for significance using the likelihood-ratio test.

Cross-validation was performed using UKB as target sample and GS:SFHS as discovery sample. Additional analyses using PRS_D_ and PRS_N_ weighted by PGC2 MDD [42] and GPC neuroticism [77] summary statistics were also tested. MDD status on UKB was adjusted by centre, array and genotyping batch as random effects and scaled (between 0 and 1) prior to analysis, giving a quasi-binomial distribution of MDD status on UKB. Models implemented on UKB (quasi-binomial regression) were adjusted by sex, age and 15 PCs. Nagelkerke’s R^2^ coefficients were estimated to quantify the proportion of MDD liability explained at the observed scale by each model and converted into R^2^ coefficients at the liability scale (prevalence: 12.2% in GS:SFHS [57] and 25.8% in UKB [63]) using Hong Lee’s transformation [79] available from GEAR: GEnetic Analysis Repository [80].

#### Using stress-sensitivity to stratify depression

GS:SFHS MDD cases (*n*_cases_ = 2,016; *n*_female_ = 1,345, *n*_male_ = 671) have data available on MDD course (single or recurrent), age of onset (*n* = 1,964) and episode count (*n* = 2,016), as well as on neuroticism (*n* = 2,010). In addition, a subset were evaluated by Mood Disorder Questionnaire [81] (MDQ; *n* = 1,022) and Schizotypal Personality Questionnaire [82] (SPQ; *n* = 1,093). The reduced sample number of MDQ and SPQ reflects the later addition of these questionnaires to the study and does not reflect a particular subgroup of GS:SFHS.

Difference in PRS_SS_ and PRS_D_ between MDD cases and controls on GS:SFHS were tested using a Student’s two sample t-test (two tailed). Cases of MDD on GS:SFHS with data available on each trait analyzed were stratified by quintiles based on PRS_SS_ and PRS_D_ (5×5 groups). Post hoc, the effects on each trait of quintiles based on PRS_SS_ and its interaction effect with quintiles based on PRS_D_ were assessed using linear regression models adjusting by sex and age in an attempt to identify a characteristic subtype of MDD patients with differential stress-sensitivity levels. The same analysis was reproduced using PRSs as continuous variables.

## Results

We confirmed the elevated neuroticism score in MDD cases in our samples. Individuals with a diagnosis of MDD had significantly higher EPQN scores compared to healthy controls (all *p* < 1.9.×10^-279^) in both GS:SFHS (mean_controls_ = 3.16; mean_cases_ = 6.42) and UKB (mean_controls_ = 2.79; mean_cases_ = 5.64). Neuroticism levels differ significantly between males and females. To control for this and any age/polygenic effects, which may account for differences in the prevalence of MDD, we created a matched set of cases and controls. The difference in neuroticism levels between cases and controls remained significant after matching the controls for PGC PRS_D_, sex and age. (GS:SFHS: mean_controls_ = 3.51; UKB: mean_controls_ = 2.97; all *p* < 2.7×10^-158^; S1 Table).

### Meta-analysis of stress-sensitivity in UKB and GS:SFHS

No SNPs were associated with stress-sensitivity at the genome-wide significant threshold (*p* < 5×10^-8^, Fig 1). However, 14 SNPs from 8 loci achieved suggestive *p* value (*p* < 1×10^-5^) ranging between *p* = 8.9×10^-6^-5.1×10^-7^ (Meta-analysis: Table 1; UKB and GS:SFHS: S2 and S3 Tables; Meta-analysis QQ-plot with *λ*: S2 Fig; UKB and GS:SFHS QQ-plots: S3 Fig). Traits with prior evidence of association with the nearest genes to the 8 lead SNPs were identified using dbGap and are shown in S4 Table. Comparison between the SNP association profile along the genome between stress-sensitivity GWIS and MDD GWAS meta-analyses is shown in Miami plots filtering for the most significant stress-sensitivity or MDD SNPs (*p* < 0.001; Meta-analysis: Fig 2; UKB and GS:SFHS: S4 Fig). No SNP with a *p*-value < 0.01 had a corresponding *p*-value in the alternate trait, suggesting that different variants contribute to depression and stress-sensitivity. Gene-based test identified *ZNF366* as the only gene achieving genome-wide significance (*p* = 1.48×10^-7^; Bonferroni-corrected significance threshold *p* < 2.79×10^-6^;S5 Table and S5 Fig). Using summary statistics from meta-analysis GWIS results, stress-sensitivity SNP-based heritability was estimated from LD score regression at 5.0% (h^2^ = 0.0499, s.e. = 0.017, *p* = 1.67×10^-3^). Conversely, the SNP-based heritability for MDD and neuroticism were estimated at 9.6% (h^2^ = 0.0962, s.e. = 0.0179, *p* = 3.87×10^-8^) and 10.1% (h^2^ = 0.1006, s.e. = 0.0076, *p* = 3.47×10^-40^) respectively, using summary statistics from the meta-analysed GWAS of UKB and GS:SFHS.

**Fig 1.**
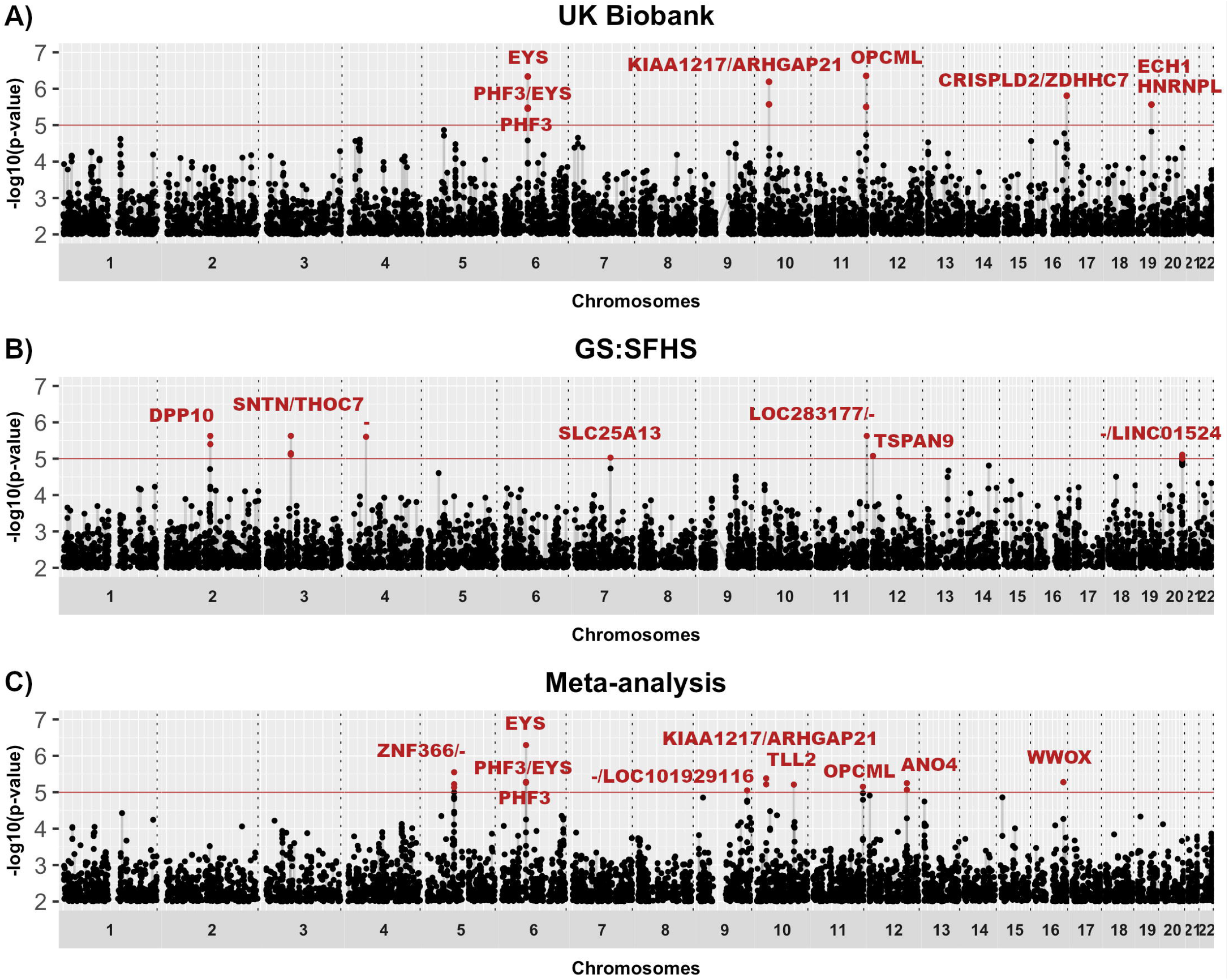
Manhattan plots showing stress-sensitivity associations. Manhattan plots of the GWIS from (A) UKB, (B) GS:SFHS and (C) sample size weighted meta-analysis of UKB and GS:SFHS. The x-axis is chromosomal position and y-axis is the *p* value (-log10 *p* value) of association with stress-sensitivity effect. Suggestive genome-wide significance threshold (*p* = 1×10^-5^) is shown by solid line at y = 5. Genes or closest gene up- and down-stream from SNP position (/) are annotated. “-”: No gene within 100kb of the SNP.

**Table 1.**
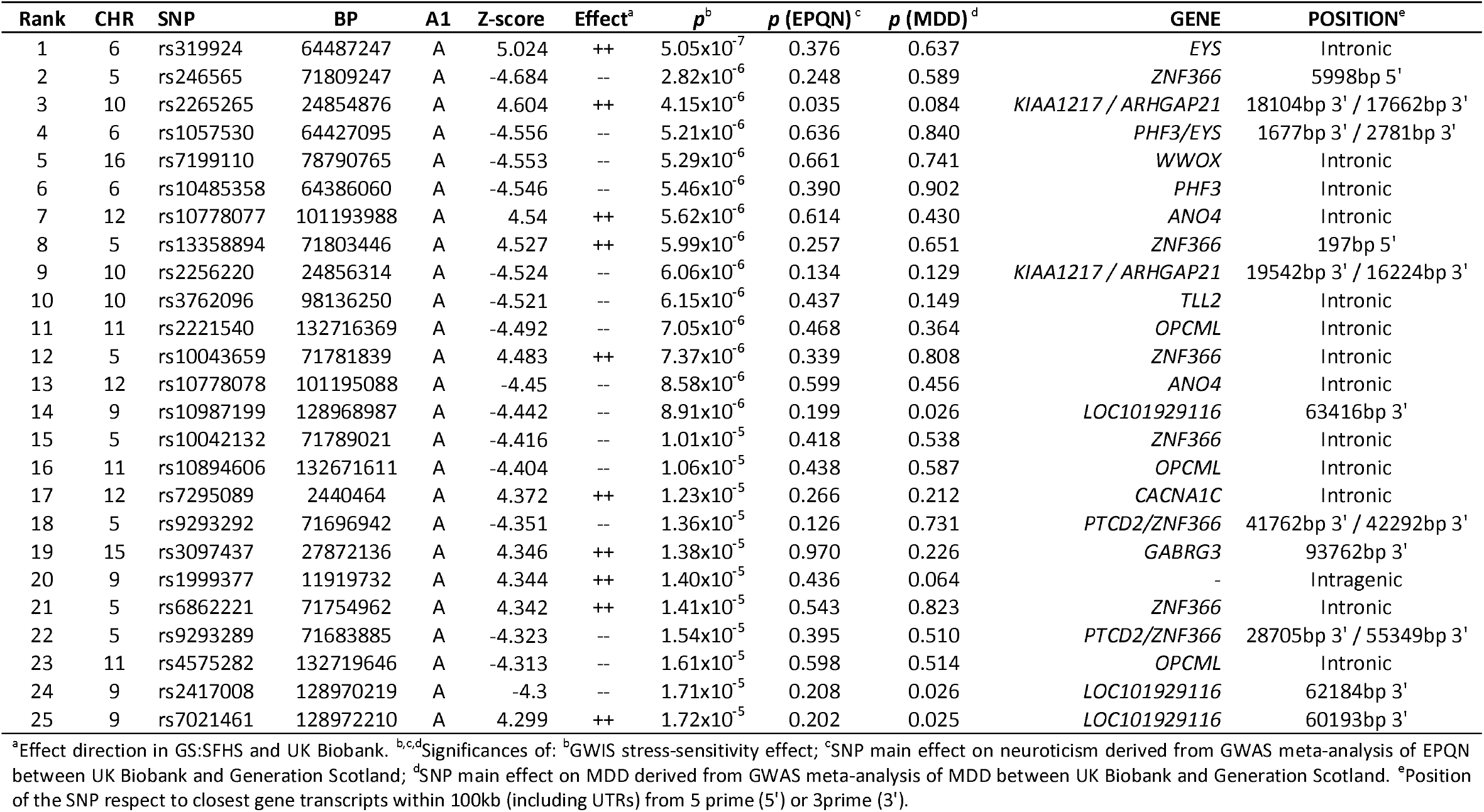
Top 25 SNPs from meta-analysis of GWISs.

**Fig 2.**
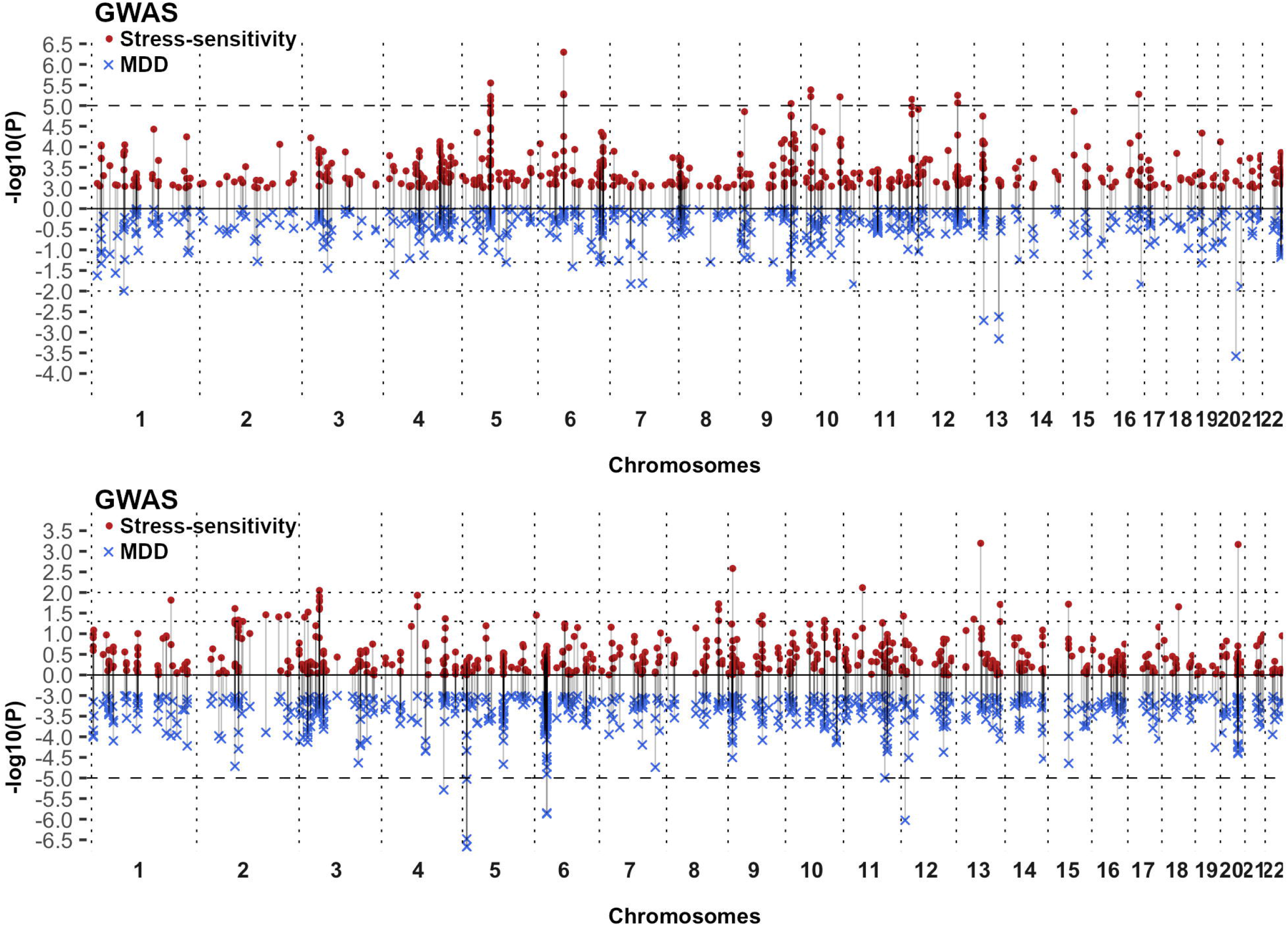
Miami plots showing comparison between association profile between stress-sensitivity GWIS and MDD GWAS. Miami plots from meta-analysis filter at *p* = 1×10^-3^: (A) filtering for stress-sensitivity *p* values (•), (B) filtering for MDD *p* values (×). The x-axis is chromosomal position and y-axis is the *p* value (-log10 *p* value) of association with stress-sensitivity (up; red dots) and MDD *p* value (down; blue crosses). Dot line: genome-wide suggestive threshold (*p* = 1×10^-5^) at the filtered effect; dashed lines: *p* = 0.01 and 0.05 at unfiltered effect.

LD score regression was performed to obtain genetic correlations between stress-sensitivity, MDD and neuroticism. As previously shown, there was a significant genetic correlation between MDD and neuroticism (r_G_ = 0.637, s.e. = 0.0704, *p* = 1.39×10^-19^). However, we found no evidence for a genetic correlation between stress-sensitivity and MDD (rG = −0.099, s.e. = 0.182, *p* = 0.585) or between stress-sensitivity and neuroticism (r_G_ = 0.114, s.e. = 0.107, *p* = 0.285).

### Pathway enrichment, functional annotation and gene expression analyses

Lead SNPs from the GWIS meta-analysis were investigated using DEPICT. No gene showed statistically significant links to stress-sensitivity at a DEPICT false discovery rate (FDR) < 0.05. No significant result was found for either gene set analysis or tissue enrichment analysis at FDR < 0.05. Evidence of regulatory elements on normal cell lines/tissues was identified for 5 of the 12 lead SNPs (i.e. rs3762096, rs10987199, rs2221540, rs246565, rs319924). Two lead SNPs were associated with eQTLs: rs319924 (an intronic SNP in *EYS*) and rs9509508 (an intronic SNP in *LATS2*) and potentially regulate *LGSN/RP3-407E4.3* (*p* = 6.31×10^-12^/*p* = 1.15×10^-5^) and *LATS2* (*p* = 3.74×10^-8^), respectively.

### Polygenic risk scores for stress-sensitivity predict MDD liability

PRS were used to investigate whether common variants affecting stress-sensitivity predict MDD risk. We generated PRS (PRS_SS_) for stress-sensitivity based on the summary statistics from the GWIS. After 10,000 permutations, PRS_SS_ significantly predicted MDD risk in GS:SFHS using weights from the larger UKB summary data (Empirical-*p* = 0.04; *p* = 5.2×10^-3^; β = 0.078, s.e. = 0.028; best-fit *p* threshold = 0.005; S6 Table). On the liability scale, the MDD variance explained in GS:SFHS by PRS_SS_ was modest (R^2^ = 0.195%). This was less than predicted by PRS weighted by the genetic main effects of MDD or neuroticism (PRS_D_: R^2^ = 0.368%; PRS_n_: R^2^ = 0.459%; Table 2 and S6 Table). However, this association was not cross-validated in UKB using summary data from the smaller GS:SFHS GWIS (Empirical-*p* = 0.68; *p* = 0.23; β = 0.004, s.e. = 0.003; best-fit *p* threshold = 0.005; PRS_SS_ R^2^ = 0.013%; S6 Table), likely due to lack of power as a result of the small discovery sample size. PRS_D_ (R^2^ = 0.204%) and PRS_N_ (R^2^ = 0.166%) derived from GS:SFHS significantly predicted MDD in UKB (Table 2 and S6 Table).

**Table 2.** MDD risk prediction at best fits.

| *UKB predicting on GS:SFHS* |  |  |  |  |  |  |
| --- | --- | --- | --- | --- | --- | --- |
| Weighted effect | Best fit threshold | # SNPs | R<sup>2</sup> (%)<sup>d</sup> | R<sup>2</sup> (%)<sup>e</sup> | *p* | Empirical-*p* |
| Stress-sensitivity | 0.005 | 1,626 | 0.141 | 0.195 | 5.2x10<sup>-3</sup> | 0.0399 |
| MDD<sup>a</sup> | 0.1 | 22,771 | 0.265 | 0.368 | 1.3x10<sup>-4</sup> | 0.0015 |
| EPQN<sup>b</sup> | 0.4 | 65,276 | 0.330 | 0.459 | 1.8x10<sup>-5</sup> | 0.0002 |
| MDD<sup>a</sup> + EPQN<sup>b</sup> | - | - | 0.503 | 0.699 | 8.0x10<sup>-7</sup> | - |
| joint models<sup>c</sup> | - | - | 0.627 | 0.871 | 1.2x10<sup>-7</sup> | - |
| *PGC2 & GPC predicting on GS:SFHS* |  |  |  |  |  |  |
| PGC2 MDD<sup>a</sup> | 1 | 92,248 | 0.993 | 1.378 | 1.4x10<sup>-13</sup> | ≤0.0001 |
| GPC EPQN<sup>b</sup> | 0.01 | 3,521 | 0.108 | 0.149 | 0.014 | 0.1038 |
| PGC2 MDD + GPC EPQN<sup>b</sup> | - | - | 1.052 | 1.461 | 1.7x10<sup>-13</sup> | - |
| joint models<sup>c</sup> | - | - | 1.203 | 1.671 | 1.6x10<sup>-14</sup> | - |
| *GS:SFHS predicting on UKB* |  |  |  |  |  |  |
| Weighted effect | Best fit threshold | # SNPs | R<sup>2</sup> (%)<sup>a</sup> | R<sup>2</sup> (%)<sup>b</sup> | *p* | Empirical-*p* |
| Stress-sensitivity | 0.005 | 1,526 | 0.008 | 0.013 | 0.231 | 0.6841 |
| MDD<sup>a</sup> | 0.03 | 7,725 | 0.130 | 0.204 | 1.6x10<sup>-6</sup> | ≤0.0001 |
| EPQN<sup>b</sup> | 0.05 | 12,296 | 0.106 | 0.166 | 1.6x10<sup>-5</sup> | 0.0005 |
| MDD<sup>a</sup> + EPQN<sup>b</sup> | - | - | 0.197 | 0.309 | 2.8x10<sup>-8</sup> | - |
| joint models<sup>c</sup> | - | - | 0.206 | 0.322 | 6.6x10<sup>-8</sup> | - |
| *PGC2 & GPC predicting on UKB* |  |  |  |  |  |  |
| PGC2 MDD<sup>a</sup> | 0.5 | 64,113 | 0.919 | 1.440 | 3.4x10<sup>-37</sup> | <0.0001 |
| GPC EPQN<sup>b</sup> | 0.03 | 8,761 | 0.066 | 0.104 | 6.5x10<sup>-4</sup> | 0.006 |
| PGC2 MDD<sup>a</sup> + GPC EPQN<sup>b</sup> | - | - | 0.950 | 1.488 | 2.9x10<sup>-37</sup> | - |
| joint models<sup>c</sup> | - | - | 0.958 | 1.501 | 1.5x10<sup>-36</sup> | - |

<sup>a</sup>major depressive disorder, <sup>b</sup>neuroticism score, <sup>c</sup>combined effect fitting all 3 PRS weighted by all the effects (i.e. stress-sensitivity, MDD and EPQN), <sup>d</sup>Nagelkerke's R<sup>2</sup> at observed scale, <sup>e</sup>R<sup>2</sup> on the liability scale.

Due to the known genetic correlations between MDD, neuroticism and stressful life events [21], models jointly fitting the effects of multiple PRS were analysed. Multiple regression analyses in GS:SFHS showed that, compared to PRS_D_ effects alone, the stress-sensitivity effect derived from the UKB GWIS effects significantly explains an additional 0.195% (a predictive improvement of 53.1%, *p* = 5.1×10^-3^; PRS_D_: β = 0.112, s.e. = 0.029; PRS_SS_ : β = 0.078, s.e. = 0.028). The inclusion of PRS_SS_ in the full model, where PRS_SS_ was fitted along with both PRS_D_ and PRS_N_ weighted by GWAS summary statistics derived from UKB remained significant; explaining an additional 0.172% (a predictive improvement of 24.6%, *p* = 8.5×10^-3^; PRS_D_: β = 0.093, s.e. = 0.029; PRS_N_: β = 0.107, s.e. = 0.030; PRS_SS_: β = 0.073, s.e. = 0.028). In models fitting PRS_D_ and PRS_N_, the variances explained were non-additive, demonstrating the partial overlap between MDD risk prediction from PRS_D_ and PRS_N_ main additive effects. This is consistent with the known genetic correlation between these two traits. An overlap was not seen between the variance explained by PRS_SS_ effect and the variance explained by PRS_D_ and/or PRS_N_. Multiple regression analyses fitting PRS_D_ and PRS_N_ derived from worldwide consortiums (Fig 3) showed that the increased sample size from GWAS used to derive PRS_D_ resulted in an increment of MDD variance explained in GS:SFHS by PRS_D_ (from 0.368% to 1. 378%). However, there was no change in the proportion of the variance explained by the PRS_SS_ in the full model (PRS_SS_ *p* = 3.5×10^-3^). These results suggest that PRS_SS_ explains a proportion of MDD risk not accounted for by PRS_D_ or PRS_N_ at current sample sizes. However, these findings were not cross-validated in UKB using PRS_SS_ derived from GS:SFHS GWIS, likely due to lack of power as a result of the small discovery sample size (S6 Fig).

**Fig 3.**
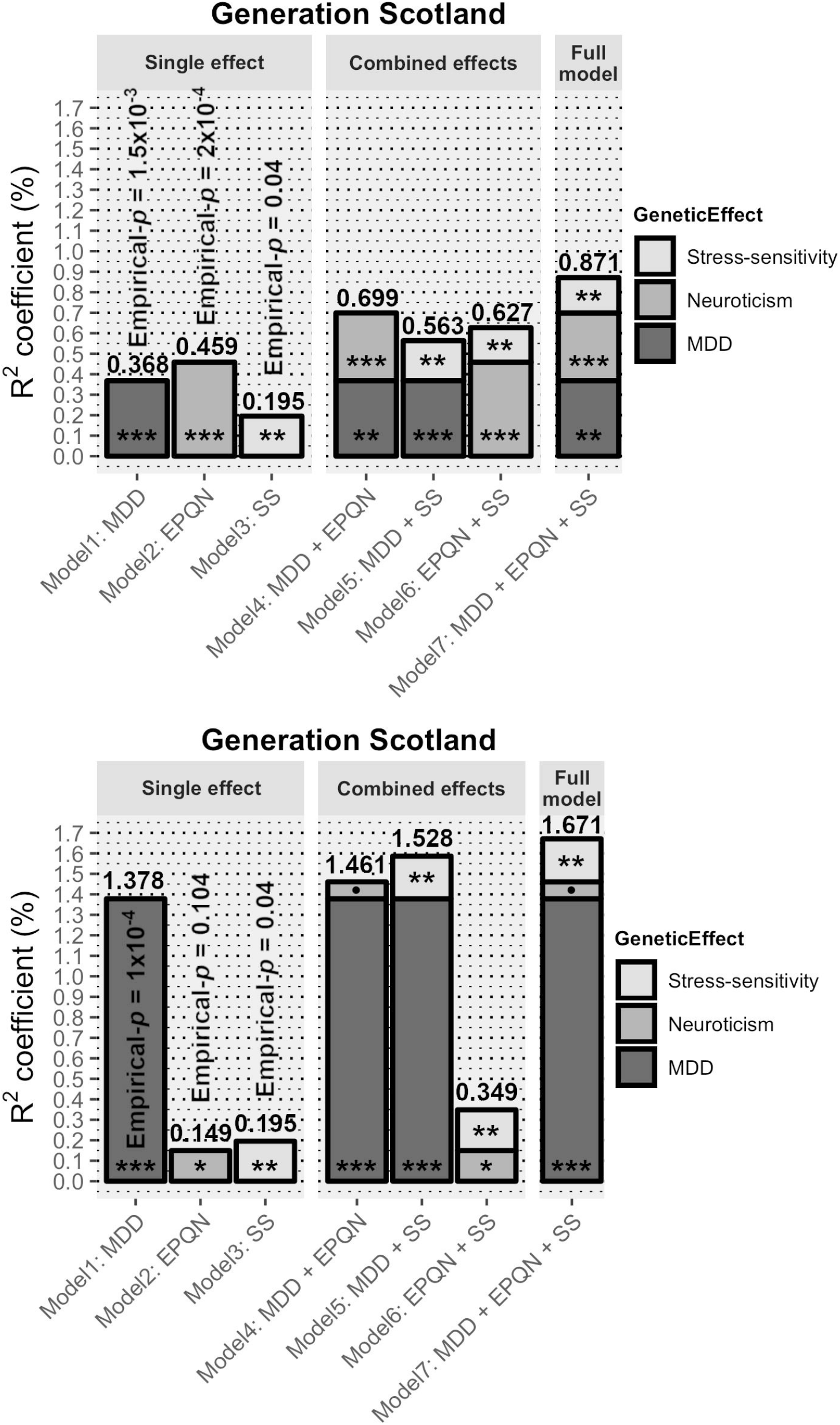
MDD is best predicted using multiple PRS. MDD risk explained (R^2^ coefficient (%); top bar values) on the liability scale by each PRS in GS:SFHS; weighted by GWAS main additive and GWIS stress-sensitivity effects independently and combined. (A) Using summary statistics from UKB as discovery sample. There is an increment on MDD risk prediction from adding PRS_SS_ to PRS_D_ model of 53.1% and 24.6% when combining PRS_SS_ with both MDD and neuroticism PRS. (B) Replication of fitting PRS_D_ and PRS_N_ using summary statistics from worldwide consortiums (i.e. PGC &GPC). Significance codes: *p* values *** < 0.001 < ** < 0.01 < * < 0.05 < • < 0.1; derived from likelihood ratio tests. *SS* stands for stress-sensitivity.

### Using stress-sensitivity to stratify MDD in GS:SFHS

MDD cases show significantly higher PRS_SS_ (*p* = 2×10^-3^) and PRS_D_ (*p* = 1.8×10^-4^) than controls. Association between MDD-related traits and stress-sensitivity risk quintiles was assessed on MDD cases in order to identify a subgroup of MDD patients, perhaps defining a characteristic aetiological subtype of MDD. However, stratification analysis failed, and no quintile based on PRS_SS_ nor its interaction with quintiles based on PRS_D_ showed statistically significant effects on any trait analyzed. Individuals with high PRS_SS_ were not significantly different from other cases for sex, MDD course, age of onset or episode count, nor neuroticism, mood disorder or schizotypal personality scores (*p* > 0.05; S7 table). Results remained non significant when PRSs were fitted as continuous variables (*p* >0.05).

## Discussion

The existence of genetic variants affecting an individual’s risk of depression in response to stress has been predicted previously [46, 49, 50] and is consistent with the departure from a simple additive genetic model seen in twin-studies of recurrent depressive disorder [83]. Through international research efforts such as the PGC and UK Biobank, there are ever-increasing sample sizes available for understanding the genetics of MDD. These resources are beginning, and will continue to, identify genome-wide significant loci [42, 84, 85]. However, the lack of environmental data and/or their reliability, makes the study of genetic individual’s response to their negative effects, and their contribution to the onset of MDD and other stress-related disorders, difficult. As a way to address this limitation, we generated a proxy for stress-sensitivity through modelling the interaction between SNP allele and MDD status on neuroticism score in a GWIS approach. Thus, we sought to identify the genetic underpinnings of individual’s sensitivity to stress response (stress-sensitivity) through those variants that contribute to higher neuroticism levels only in individuals with a lifetime diagnosis of MDD but not in healthy controls.

We performed a GWIS to identify loci showing differential effects on neuroticism scores in individuals with and without MDD (so called stress-sensitivity proxy). No SNPs reached genome-wide significance, but 14 SNPs from 8 loci reached suggestive significance levels (see S4 Table for prior evidence of associated phenotypes). Enrichment analysis showed no evidence for enrichment of specific pathways or tissues. The top two loci, *PTP4A1-PHF3-EYS* and *ZNF366* have been previously associated with alcohol dependence [86-90], alcohol intake (dbGaP: phs000342) and glucocorticoid receptor function [91-93]. The most significant SNP in this study, rs319924, is an intronic variant in *EYS* that is a potential eQTL for *LGSN* [76], a gene previously associated with male-specific depression [94]. This is of particular interest given previous studies linking alcohol consumption, stress and the risk of depression [95-100]. However, findings should be interpreted with caution, as these loci did not reach genome-wide significance at current sample size. Evidence of an eQTL effect was predicted for a lead SNP in *LATS2*, a positive regulator of histone methyltransferase activity [101] a process important in anxiety-related behaviours [102]. The prior association of the top two loci in this study with alcohol related-phenotypes suggests that genes involved in the sensitivity to stress may mediate the effects of stress on alcohol consumption. Some *PHF3* paralogs have been shown to be linked with depression and modulate stress response [103,104].

Gene-based analysis identified a genome-wide significant association between *ZNF366* and stress-sensitivity. *ZNF366* (also known as *DC-SCRIPT*) is a corepressor of transcription found in nuclear receptor complexes including the glucocorticoid receptor. *ZNF366* represses glucocorticoid receptor-mediated transcription in monocyte-derived dendritic cells [91]; and may act through histone deacetylases to modulate immune response [92]. There is evidence from a large-scale mRNA display study that *PHF3*, in the region underlying the most significant peak in the single SNP analysis, may also interact, directly or indirectly, with the glucocorticoid receptor (IntAct database [93]) but this has not been confirmed. These results reinforce the hypothesis that our proxy for stress-sensitivity truly reflects the genetic architecture of sensitivity to respond to stress. We estimated a significant lower bound on common SNP-based heritability for stress-sensitivity of 5%. Whilst the known genetic overlap between MDD and neuroticism was detectable, the lack of genetic correlation with stress-sensitivity, reinforced by results from multiple regression analyses, indicated a lack of significant overlap in the genetics factors underpinning stress-sensitivity and MDD or neuroticism. This analysis may be limited by our sample size, although using the largest available meta-analyses of MDD and neuroticism [42, 77] did not decrease the proportion of liability explained by the PRS_SS_. We note, that as such meta-analyses increase in size it is likely, as with the effects of smoking in schizophrenia [105, 106], that the indirect genetic effects of the environment on the risk of depression will be detected by GWAS. However, through studies such as ours, or similar, the mechanism for the effect of the risk alleles may be clarified.

Further, we show that such genetic information in stress-sensitivity could significantly improve the proportion of liability to MDD predicted by PRS based only on additive genetic effects on MDD identified by large GWAS. The summary results from the GWIS were used to derive a PRS reflecting the genetic difference in stress-sensitivity. This variable significantly predicted liability to MDD in GS:SFHS (*p* = 5.2×10^-3^, Empirical-*p* = 0.04 after 10,000 permutations), although this finding could not be replicated in UKB (Empirical-*p* = 0.68), likely due to lack of power. This is consistent with the expectation that the larger the discovery sample (i.e. UKB), the greater the accuracy of the weighting and the more predictive the PRS [107]. Multiple regression models in GS:SFHS suggest that inclusion of PRS weighted by stress-sensitivity significantly improves MDD prediction over use of either MDD and/or neuroticism weighted PRS alone (improvement in full model *p* = 8.5×10^-3^). However, we were unable to identify a subgroup of MDD cases with higher PRS_SS_. The polygenic interaction approach used in our study may, therefore, improve the interpretation of both positive and negative findings from GWAS studies (i.e. pathways and mechanisms involved, lack of replication, or negative findings in variants mediating environmental effects). Added to paralleling recent developments in GWAS analyses, it may maximize our power to detect gene-by-environment effects in this heterogeneous disorder.

Future studies will be required to further investigate the effects of adverse life events in individuals with high or low polygenic risk scores for stress-sensitivity. However, the methodology presented allows addressing the genetic response to negative outcomes via proxy in the absence of prospective environmental data.

Here we identify an independent set of risk variants for an individual’s response to negative outcomes and show that incorporating information across many loci provides clear and replicable evidence for a genetic effect of stress-sensitivity on MDD risk; identifying a potential genetic link with alcohol intake. These results require further study, but may inform treatment of comorbid alcohol dependency and depression.

## Acknowledgments

We thank all participants from UK Biobank, including families, volunteers, technicians and scientist who helped collect the samples and prepare the data. Analyses were performed under UK Biobank project 4844. We are grateful to all the families who took part in Generation Scotland, the general practitioners and the Scottish School of Primary Care for their help in recruiting them, the whole Generation Scotland team, which includes interviewers, computer and laboratory technicians, clerical workers, research scientists, volunteers, managers, receptionists, healthcare assistants and nurses. We acknowledge the Genetics Core Laboratory at the Wellcome Trust Clinical Research Facility, Edinburgh to carry out the genotyping of the Generation Scotland samples. We gratefully acknowledge **The Major Depressive Disorder Working Group of the Psychiatric Genomics Consortium,** which depends on the contributions of many parties: Naomi R Wray (Institute for Molecular Bioscience, The University of Queensland, Brisbane, QLD, AU and Queensland Brain Institute, The University of Queensland, Brisbane, QLD, AU), Stephan Ripke (Analytic and Translational Genetics Unit, Massachusetts General Hospital, Boston, MA, US, Department of Psychiatry and Psychotherapy, Universitätsmedizin Berlin Campus Charité Mitte, Berlin, DE and Medical and Population Genetics, Broad Institute, Cambridge, MA, US), Manuel Mattheisen (Centre for Psychiatry Research, Department of Clinical Neuroscience, Karolinska Institutet, Stockholm, SE, Department of Biomedicine, Aarhus University, Aarhus, DK, iSEQ, Centre for Integrative Sequencing, Aarhus University, Aarhus, DK and iPSYCH, The Lundbeck Foundation Initiative for Integrative Psychiatric Research, DK), Maciej Trzaskowski (Institute for Molecular Bioscience, The University of Queensland, Brisbane, QLD), Enda M Byrne (Institute for Molecular Bioscience, The University of Queensland, Brisbane, QLD, AU), Abdel Abdellaoui (Dept of Biological Psychology & EMGO+ Institute for Health and Care Research, Vrije Universiteit Amsterdam, Amsterdam, NL), Mark J Adams (Division of Psychiatry, University of Edinburgh, Edinburgh, GB), Esben Agerbo (iPSYCH, The Lundbeck Foundation Initiative for Integrative Psychiatric Research, DK, Centre for Integrated Register-based Research, Aarhus University, Aarhus, DK and National Centre for Register-Based Research, Aarhus University, Aarhus, DK), Tracy M Air (Discipline of Psychiatry, University of Adelaide, Adelaide, SA, AU), Till F M Andlauer (Department of Translational Research in Psychiatry, Max Planck Institute of Psychiatry, Munich, DE and Munich Cluster for Systems Neurology (SyNergy), Munich, DE), Silviu-Alin Bacanu (Department of Psychiatry, Virginia Commonwealth University, Richmond, VA, US), Marie Bækvad-Hansen (iPSYCH, The Lundbeck Foundation Initiative for Integrative Psychiatric Research, DK and Center for Neonatal Screening, Department for Congenital Disorders, Statens Serum Institut, Copenhagen, DK), Aartjan T F Beekman (Department of Psychiatry, Vrije Universiteit Medical Center and GGZ inGeest, Amsterdam, NL), Tim B Bigdeli (Department of Psychiatry, Virginia Commonwealth University, Richmond, VA, US and Virginia Institute for Psychiatric and Behavior Genetics, Richmond, VA, US), Elisabeth B Binder (Department of Translational Research in Psychiatry, Max Planck Institute of Psychiatry, Munich, DE and Department of Psychiatry and Behavioral Sciences, Emory University School of Medicine, Atlanta, GA, US), Douglas H R Blackwood (Division of Psychiatry, University of Edinburgh, Edinburgh, GB), Julien Bryois (Department of Medical Epidemiology and Biostatistics, Karolinska Institutet, Stockholm, SE), Henriette N Buttenschøn (iSEQ, Centre for Integrative Sequencing, Aarhus University, Aarhus, DK, iPSYCH, The Lundbeck Foundation Initiative for Integrative Psychiatric Research, DK and Department of Clinical Medicine, Translational Neuropsychiatry Unit, Aarhus University, Aarhus, DK), Jonas Bybjerg-Grauholm (iPSYCH, The Lundbeck Foundation Initiative for Integrative Psychiatric Research, DK and Center for Neonatal Screening, Department for Congenital Disorders, Statens Serum Institut, Copenhagen, DK), Na Cai (Human Genetics, Wellcome Trust Sanger Institute, Cambridge, GB and Statistical genomics and systems genetics, European Bioinformatics Institute (EMBL-EBI), Cambridge, GB), Enrique Castelao (Department of Psychiatry, University Hospital of Lausanne, Prilly, Vaud, CH), Jane Hvarregaard Christensen (Department of Biomedicine, Aarhus University, Aarhus, DK, iSEQ, Centre for Integrative Sequencing, Aarhus University, Aarhus, DK and iPSYCH, The Lundbeck Foundation Initiative for Integrative Psychiatric Research, DK), Toni-Kim Clarke (Division of Psychiatry, University of Edinburgh, Edinburgh, GB), Jonathan R I Coleman (MRC Social Genetic and Developmental Psychiatry Centre, King’s College London, London, GB), Lucía Colodro-Conde (Genetics and Computational Biology, QIMR Berghofer Medical Research Institute, Herston, QLD, AU), Baptiste Couvy-Duchesne (Centre for Advanced Imaging, The University of Queensland, Saint Lucia, QLD, AU and Queensland Brain Institute, The University of Queensland, Saint Lucia, QLD, AU), Nick Craddock (Psychological Medicine, Cardiff University, Cardiff, GB), Gregory E Crawford (Center for Genomic and Computational Biology, Duke University, Durham, NC, US and Department of Pediatrics, Division of Medical Genetics, Duke University, Durham, NC, US), Gail Davies (Centre for Cognitive Ageing and Cognitive Epidemiology, University of Edinburgh, Edinburgh, GB), Ian J Deary (Centre for Cognitive Ageing and Cognitive Epidemiology, University of Edinburgh, Edinburgh, GB), Franziska Degenhardt (Institute of Human Genetics, University of Bonn, Bonn, DE and Life&Brain Center, Department of Genomics, University of Bonn, Bonn, DE), Eske M Derks (Genetics and Computational Biology, QIMR Berghofer Medical Research Institute, Herston, QLD, AU), Nese Direk (Epidemiology, Erasmus MC, Rotterdam, Zuid-Holland, NL and Psychiatry, Dokuz Eylul University School Of Medicine, Izmir, TR), Conor V Dolan (Dept of Biological Psychology & EMGO+ Institute for Health and Care Research, Vrije Universiteit Amsterdam, Amsterdam, NL), Erin C Dunn (Department of Psychiatry, Massachusetts General Hospital, Boston, MA, US, Psychiatric and Neurodevelopmental Genetics Unit (PNGU), Massachusetts General Hospital, Boston, MA, US and Stanley Center for Psychiatric Research, Broad Institute, Cambridge, MA, US), Thalia C Eley (MRC Social Genetic and Developmental Psychiatry Centre, King’s College London, London, GB), Valentina Escott-Price (Neuroscience and Mental Health, Cardiff University, Cardiff, GB), Farnush Farhadi Hassan Kiadeh (Bioinformatics, University of British Columbia, Vancouver, BC, CA), Hilary K Finucane (Department of Epidemiology, Harvard T.H. Chan School of Public Health, Boston, MA, US and Department of Mathematics, Massachusetts Institute of Technology, Cambridge, MA, US), Andreas J Forstner (Institute of Human Genetics, University of Bonn, Bonn, DE, Life&Brain Center, Department of Genomics, University of Bonn, Bonn, DE, Department of Psychiatry (UPK), University of Basel, Basel, CH and Human Genomics Research Group, Department of Biomedicine, University of Basel, Basel, CH), Josef Frank (Department of Genetic Epidemiology in Psychiatry, Central Institute of Mental Health, Medical Faculty Mannheim, Heidelberg University, Mannheim, Baden-Württemberg, DE), Héléna A Gaspar (MRC Social Genetic and Developmental Psychiatry Centre, King’s College London, London, GB), Michael Gill (Department of Psychiatry, Trinity College Dublin, Dublin, IE), Fernando S Goes (Psychiatry & Behavioral Sciences, Johns Hopkins University, Baltimore, MD, US), Scott D Gordon (Genetics and Computational Biology, QIMR Berghofer Medical Research Institute, Brisbane, QLD, AU), Jakob Grove (Department of Biomedicine, Aarhus University, Aarhus, DK, iSEQ, Centre for Integrative Sequencing, Aarhus University, Aarhus, DK, iPSYCH, The Lundbeck Foundation Initiative for Integrative Psychiatric Research, DK and Bioinformatics Research Centre, Aarhus University, Aarhus, DK), Lynsey S Hall (Division of Psychiatry, University of Edinburgh, Edinburgh, GB and Institute of Genetic Medicine, Newcastle University, Newcastle upon Tyne, GB), Christine Søholm Hansen (iPSYCH, The Lundbeck Foundation Initiative for Integrative Psychiatric Research, DK and Center for Neonatal Screening, Department for Congenital Disorders, Statens Serum Institut, Copenhagen, DK), Thomas F Hansen (Danish Headache Centre, Department of Neurology, Rigshospitalet, Glostrup, DK, Institute of Biological Psychiatry, Mental Health Center Sct. Hans, Mental Health Services Capital Region of Denmark, Copenhagen, DK and iPSYCH, The Lundbeck Foundation Initiative for Psychiatric Research, Copenhagen, DK), Stefan Herms (Institute of Human Genetics, University of Bonn, Bonn, DE, Life&Brain Center, Department of Genomics, University of Bonn, Bonn, DE and Human Genomics Research Group, Department of Biomedicine, University of Basel, Basel, CH), Ian B Hickie (Brain and Mind Centre, University of Sydney, Sydney, NSW, AU), Per Hoffmann (Institute of Human Genetics, University of Bonn, Bonn, DE, Life&Brain Center, Department of Genomics, University of Bonn, Bonn, DE and Human Genomics Research Group, Department of Biomedicine, University of Basel, Basel, CH), Georg Homuth (Interfaculty Institute for Genetics and Functional Genomics, Department of Functional Genomics, University Medicine and Ernst Moritz Arndt University Greifswald, Greifswald, Mecklenburg-Vorpommern, DE), Carsten Horn (Roche Pharmaceutical Research and Early Development, Pharmaceutical Sciences, Roche Innovation Center Basel, F. Hoffmann-La Roche Ltd, Basel, CH), Jouke-Jan Hottenga (Dept of Biological Psychology & EMGO+ Institute for Health and Care Research, Vrije Universiteit Amsterdam, Amsterdam, NL), David M Hougaard (iPSYCH, The Lundbeck Foundation Initiative for Integrative Psychiatric Research, DK and Center for Neonatal Screening, Department for Congenital Disorders, Statens Serum Institut, Copenhagen, DK), Marcus Ising (Max Planck Institute of Psychiatry, Munich, DE), Rick Jansen (Department of Psychiatry, Vrije Universiteit Medical Center and GGZ inGeest, Amsterdam, NL), Eric Jorgenson (Division of Research, Kaiser Permanente Northern California, Oakland, CA, US), James A Knowles (Psychiatry & The Behavioral Sciences, University of Southern California, Los Angeles, CA, US), Isaac S Kohane (Department of Biomedical Informatics, Harvard Medical School, Boston, MA, US, Department of Medicine, Brigham and Women’s Hospital, Boston, MA, US and Informatics Program, Boston Children’s Hospital, Boston, MA, US), Julia Kraft (Department of Psychiatry and Psychotherapy, Universitätsmedizin Berlin Campus Charité Mitte, Berlin, DE), Warren W. Kretzschmar (Wellcome Trust Centre for Human Genetics, University of Oxford, Oxford, GB), Jesper Krogh (Department of Endocrinology at Herlev University Hospital, University of Copenhagen, Copenhagen, DK), Zoltán Kutalik (Institute of Social and Preventive Medicine (IUMSP), University Hospital of Lausanne, Lausanne, VD, CH and Swiss Institute of Bioinformatics, Lausanne, VD, CH), Yihan Li (Wellcome Trust Centre for Human Genetics, University of Oxford, Oxford, GB), Penelope A Lind (Genetics and Computational Biology, QIMR Berghofer Medical Research Institute, Herston, QLD, AU), Donald J MacIntyre Division of Psychiatry, Centre for Clinical Brain Sciences, University of Edinburgh, Edinburgh, GB and Mental Health, NHS 24, Glasgow, GB), Dean F MacKinnon (Psychiatry & Behavioral Sciences, Johns Hopkins University, Baltimore, MD, US), Robert M Maier (Queensland Brain Institute, The University of Queensland, Brisbane, QLD, AU), Wolfgang Maier (Department of Psychiatry and Psychotherapy, University of Bonn, Bonn, DE), Jonathan Marchini (Statistics, University of Oxford, Oxford, GB), Hamdi Mbarek (Dept of Biological Psychology & EMGO+ Institute for Health and Care Research, Vrije Universiteit Amsterdam, Amsterdam, NL), Patrick McGrath (Psychiatry, Columbia University College of Physicians and Surgeons, New York, NY, US), Peter McGuffin (MRC Social Genetic and Developmental Psychiatry Centre, King’s College London, London, GB), Sarah E Medland (Genetics and Computational Biology, QIMR Berghofer Medical Research Institute, Herston, QLD, AU), Divya Mehta (Queensland Brain Institute, The University of Queensland, Brisbane, QLD, AU and School of Psychology and Counseling, Queensland University of Technology, Brisbane, QLD, AU), Christel M Middeldorp (Dept of Biological Psychology & EMGO+ Institute for Health and Care Research, Vrije Universiteit Amsterdam, Amsterdam, NL, Child and Youth Mental Health Service, Children’s Health Queensland Hospital and Health Service, South Brisbane, QLD, AU and Child Health Research Centre, University of Queensland, Brisbane, QLD, AU), Evelin Mihailov (Estonian Genome Center, University of Tartu, Tartu, EE), Yuri Milaneschi (Department of Psychiatry, Vrije Universiteit Medical Center and GGZ inGeest, Amsterdam, NL), Lili Milani (Estonian Genome Center, University of Tartu, Tartu, EE), Francis M Mondimore (Psychiatry & Behavioral Sciences, Johns Hopkins University, Baltimore, MD, US), Grant W Montgomery (Institute for Molecular Bioscience, The University of Queensland, Brisbane, QLD, AU), Sara Mostafavi (Medical Genetics, University of British Columbia, Vancouver, BC, CA and Statistics, University of British Columbia, Vancouver, BC, CA), Niamh Mullins (MRC Social Genetic and Developmental Psychiatry Centre, King’s College London, London, GB), Matthias Nauck DZHK (German Centre for Cardiovascular Research), Partner Site Greifswald, University Medicine, University Medicine Greifswald, Greifswald, Mecklenburg-Vorpommern, DE and Institute of Clinical Chemistry and Laboratory Medicine, University Medicine Greifswald, Greifswald, Mecklenburg-Vorpommern, DE), Bernard Ng (Statistics, University of British Columbia, Vancouver, BC, CA), Michel G Nivard (Dept of Biological Psychology & EMGO+ Institute for Health and Care Research, Vrije Universiteit Amsterdam, Amsterdam, NL), Dale R Nyholt (Institute of Health and Biomedical Innovation, Queensland University of Technology, Brisbane, QLD, AU), Paul F O’Reilly (MRC Social Genetic and Developmental Psychiatry Centre, King’s College London, London, GB), Hogni Oskarsson (Humus, Reykjavik, IS), Michael J Owen (MRC Centre for Neuropsychiatric Genetics and Genomics, Cardiff University, Cardiff, GB), Jodie N Painter (Genetics and Computational Biology, QIMR Berghofer Medical Research Institute, Herston, QLD, AU), Carsten Bøcker Pedersen (iPSYCH, The Lundbeck Foundation Initiative for Integrative Psychiatric Research, DK, Centre for Integrated Register-based Research, Aarhus University, Aarhus, DK and National Centre for Register-Based Research, Aarhus University, Aarhus, DK), Marianne Giørtz Pedersen (iPSYCH, The Lundbeck Foundation Initiative for Integrative Psychiatric Research, DK, Centre for Integrated Register-based Research, Aarhus University, Aarhus, DK and National Centre for Register-Based Research, Aarhus University, Aarhus, DK), Roseann E. Peterson (Department of Psychiatry, Virginia Commonwealth University, Richmond, VA, US and Virginia Institute for Psychiatric & Behavioral Genetics, Virginia Commonwealth University, Richmond, VA, US), Erik Pettersson (Department of Medical Epidemiology and Biostatistics, Karolinska Institutet, Stockholm, SE), Wouter J Peyrot (Department of Psychiatry, Vrije Universiteit Medical Center and GGZ inGeest, Amsterdam, NL), Giorgio Pistis (Department of Psychiatry, University Hospital of Lausanne, Prilly, Vaud, CH), Danielle Posthuma (Clinical Genetics, Vrije Universiteit Medical Center, Amsterdam, NL and Complex Trait Genetics, Vrije Universiteit Amsterdam, Amsterdam, NL), Jorge A Quiroz (Solid Biosciences, Boston, MA, US), Per Qvist (Department of Biomedicine, Aarhus University, Aarhus, DK, iSEQ, Centre for Integrative Sequencing, Aarhus University, Aarhus, DK and iPSYCH, The Lundbeck Foundation Initiative for Integrative Psychiatric Research, DK), John P Rice (Department of Psychiatry, Washington University in Saint Louis School of Medicine, Saint Louis, MO, US), Brien P. Riley (Department of Psychiatry, Virginia Commonwealth University, Richmond, VA, US), Margarita Rivera (MRC Social Genetic and Developmental Psychiatry Centre, King’s College London, London, GB and Department of Biochemistry and Molecular Biology II, Institute of Neurosciences, Center for Biomedical Research, University of Granada, Granada, ES), Saira Saeed Mirza (Epidemiology, Erasmus MC, Rotterdam, Zuid-Holland, NL), Robert Schoevers (Department of Psychiatry, University of Groningen, University Medical Center Groningen, Groningen, NL), Eva C Schulte (Department of Psychiatry and Psychotherapy, Medical Center of the University of Munich, Campus Innenstadt, Munich, DE and Institute of Psychiatric Phenomics and Genomics (IPPG), Medical Center of the University of Munich, Campus Innenstadt, Munich, DE), Ling Shen (Division of Research, Kaiser Permanente Northern California, Oakland, CA, US), Jianxin Shi (Division of Cancer Epidemiology and Genetics, National Cancer Institute, Bethesda, MD, US), Stanley I Shyn (Behavioral Health Services, Kaiser Permanente Washington, Seattle, WA, US), Engilbert Sigurdsson (Faculty of Medicine, Department of Psychiatry, University of Iceland, Reykjavik, IS), Grant C B Sinnamon (School of Medicine and Dentistry, James Cook University, Townsville, QLD, AU), Johannes H Smit (Department of Psychiatry, Vrije Universiteit Medical Center and GGZ inGeest, Amsterdam, NL), Daniel J Smith (Institute of Health and Wellbeing, University of Glasgow, Glasgow, GB), Hreinn Stefansson (deCODE Genetics / Amgen, Reykjavik, IS), Stacy Steinberg (deCODE Genetics / Amgen, Reykjavik, IS), Fabian Streit (Department of Genetic Epidemiology in Psychiatry, Central Institute of Mental Health, Medical Faculty Mannheim, Heidelberg University, Mannheim, Baden-Württemberg, DE), Jana Strohmaier (Department of Genetic Epidemiology in Psychiatry, Central Institute of Mental Health, Medical Faculty Mannheim, Heidelberg University, Mannheim, Baden-Württemberg, DE), Katherine E Tansey (College of Biomedical and Life Sciences, Cardiff University, Cardiff, GB), Henning Teismann (Institute of Epidemiology and Social Medicine, University of Münster, Münster, Nordrhein-Westfalen, DE), Alexander Teumer (Institute for Community Medicine, University Medicine Greifswald, Greifswald, Mecklenburg-Vorpommern, DE), Wesley Thompson iPSYCH, The Lundbeck Foundation Initiative for Integrative Psychiatric Research, DK, Institute of Biological Psychiatry, Mental Health Center Sct. Hans, Mental Health Services Capital Region of Denmark, Copenhagen, DK, Department of Psychiatry, University of California, San Diego, San Diego, CA, US and KG Jebsen Centre for Psychosis Research, Norway Division of Mental Health and Addiction, Oslo University Hospital, Oslo, NO), Pippa A Thomson (Medical Genetics Section, CGEM, IGMM, University of Edinburgh, Edinburgh, GB), Thorgeir E Thorgeirsson (deCODE Genetics / Amgen, Reykjavik, IS), Matthew Traylor (Clinical Neurosciences, University of Cambridge, Cambridge, GB), Jens Treutlein (Department of Genetic Epidemiology in Psychiatry, Central Institute of Mental Health, Medical Faculty Mannheim, Heidelberg University, Mannheim, Baden-Württemberg, DE), Vassily Trubetskoy (Department of Psychiatry and Psychotherapy, Universitätsmedizin Berlin Campus Charité Mitte, Berlin, DE), André G Uitterlinden (Internal Medicine, Erasmus MC, Rotterdam, Zuid-Holland, NL), Daniel Umbricht (Roche Pharmaceutical Research and Early Development, Neuroscience, Ophthalmology and Rare Diseases Discovery & Translational Medicine Area, Roche Innovation Center Basel, F. Hoffmann-La Roche Ltd, Basel, CH), Sandra Van der Auwera (Department of Psychiatry and Psychotherapy, University Medicine Greifswald, Greifswald, Mecklenburg-Vorpommern, DE), Albert M van Hemert (Department of Psychiatry, Leiden University Medical Center, Leiden, NL), Alexander Viktorin (Department of Medical Epidemiology and Biostatistics, Karolinska Institutet Stockholm, SE), Peter M Visscher (Institute for Molecular Bioscience, The University of Queensland, Brisbane, QLD, AU and Queensland Brain Institute, The University of Queensland, Brisbane, QLD, AU), Yunpeng Wang (iPSYCH, The Lundbeck Foundation Initiative for Integrative Psychiatric Research, DK, Institute of Biological Psychiatry, Mental Health Center Sct. Hans, Mental Health Services Capital Region of Denmark, Copenhagen, DK and KG Jebsen Centre for Psychosis Research, Norway Division of Mental Health and Addiction, Oslo University Hospital, Oslo, NO), Bradley T. Webb (Virginia Institute of Psychiatric & Behavioral Genetics, Virginia Commonwealth University, Richmond, VA, US), Shantel Marie Weinsheimer (iPSYCH, The Lundbeck Foundation Initiative for Integrative Psychiatric Research, DK and Institute of Biological Psychiatry, Mental Health Center Sct. Hans, Mental Health Services Capital Region of Denmark, Copenhagen, DK), Jürgen Wellmann (Institute of Epidemiology and Social Medicine, University of Münster, Münster, Nordrhein-Westfalen, DE), Gonneke Willemsen (Dept of Biological Psychology & EMGO+ Institute for Health and Care Research, Vrije Universiteit Amsterdam, Amsterdam, NL), Stephanie H Witt (Department of Genetic Epidemiology in Psychiatry, Central Institute of Mental Health, Medical Faculty Mannheim, Heidelberg University, Mannheim, Baden-Württemberg, DE), Yang Wu (Institute for Molecular Bioscience, The University of Queensland, Brisbane, QLD, AU), Hualin S Xi (Computational Sciences Center of Emphasis, Pfizer Global Research and Development, Cambridge, MA, US), Jian Yang (Queensland Brain Institute, The University of Queensland, Brisbane, QLD, AU and Institute for Molecular Bioscience; Queensland Brain Institute, The University of Queensland, Brisbane, QLD, AU), Futao Zhang (Institute for Molecular Bioscience, The University of Queensland, Brisbane, QLD, AU), Volker Arolt (Department of Psychiatry, University of Münster, Münster, Nordrhein-Westfalen, DE), Bernhard T Baune (Discipline of Psychiatry, University of Adelaide, Adelaide, SA, AU), Klaus Berger (Institute of Epidemiology and Social Medicine, University of Münster, Münster, Nordrhein-Westfalen, DE), Dorret I Boomsma (Dept of Biological Psychology & EMGO+ Institute for Health and Care Research, Vrije Universiteit Amsterdam, Amsterdam, NL), Sven Cichon (Institute of Human Genetics, University of Bonn, Bonn, DE, Human Genomics Research Group, Department of Biomedicine, University of Basel, Basel, CH, Institute of Medical Genetics and Pathology, University Hospital Basel, University of Basel, Basel, CH and Institute of Neuroscience and Medicine (INM-1), Research Center Juelich, Juelich, DE), Udo Dannlowski (Department of Psychiatry, University of Münster, Münster, Nordrhein-Westfalen, DE), EJC de Geus (Dept of Biological Psychology & EMGO+ Institute for Health and Care Research, Vrije Universiteit Amsterdam, Amsterdam, NL and Amsterdam Public Health Institute, Vrije Universiteit Medical Center, Amsterdam, NL), J Raymond DePaulo (Psychiatry & Behavioral Sciences, Johns Hopkins University, Baltimore, MD, US), Enrico Domenici (Centre for Integrative Biology, Università degli Studi di Trento, Trento, Trentino-Alto Adige, IT), Katharina Domschke (Department of Psychiatry and Psychotherapy, Medical Center, University of Freiburg, Faculty of Medicine, University of Freiburg, Freiburg, DE), Tõnu Esko 5, (Estonian Genome Center, University of Tartu, Tartu, EE), Hans J Grabe (Department of Psychiatry and Psychotherapy, University Medicine Greifswald, Greifswald, Mecklenburg-Vorpommern, DE), Steven P Hamilton (Psychiatry, Kaiser Permanente Northern California, San Francisco, CA, US), Caroline Hayward (Medical Research Council Human Genetics Unit, Institute of Genetics and Molecular Medicine, University of Edinburgh, Edinburgh, GB), Andrew C Heath (Department of Psychiatry, Washington University in Saint Louis School of Medicine, Saint Louis, MO, US), Kenneth S Kendler (Department of Psychiatry, Virginia Commonwealth University, Richmond, VA, US), Stefan Kloiber (Max Planck Institute of Psychiatry, Munich, DE, Department of Psychiatry, University of Toronto, Toronto, ON, CA and Centre for Addiction and Mental Health, Toronto, ON, CA), Glyn Lewis (Division of Psychiatry, University College London, London, GB), Qingqin S Li (Neuroscience Therapeutic Area, Janssen Research and Development, LLC, Titusville, NJ, US), Susanne Lucae (Max Planck Institute of Psychiatry, Munich, DE), Pamela AF Madden (Department of Psychiatry, Washington University in Saint Louis School of Medicine, Saint Louis, MO, US), Patrik K Magnusson (Department of Medical Epidemiology and Biostatistics, Karolinska Institutet, Stockholm, SE), Nicholas G Martin (Genetics and Computational Biology, QIMR Berghofer Medical Research Institute, Brisbane, QLD, AU), Andrew M McIntosh* (Division of Psychiatry, University of Edinburgh, Edinburgh, GB and Centre for Cognitive Ageing and Cognitive Epidemiology, University of Edinburgh, Edinburgh, GB), Andres Metspalu (Estonian Genome Center, University of Tartu, Tartu, EE and Institute of Molecular and Cell Biology, University of Tartu, Tartu, EE), Ole Mors (iPSYCH, The Lundbeck Foundation Initiative for Integrative Psychiatric Research, DK and Psychosis Research Unit, Aarhus University Hospital, Risskov, Aarhus, DK), Preben Bo Mortensen (iSEQ, Centre for Integrative Sequencing, Aarhus University, Aarhus, DK, iPSYCH, The Lundbeck Foundation Initiative for Integrative Psychiatric Research, DK, Centre for Integrated Register-based Research, Aarhus University, Aarhus, DK and National Centre for Register-Based Research, Aarhus University, Aarhus, DK), Bertram Müller-Myhsok (Department of Translational Research in Psychiatry, Max Planck Institute of Psychiatry, Munich, DE, Munich Cluster for Systems Neurology (SyNergy), Munich, DE and University of Liverpool, Liverpool, GB), Merete Nordentoft (iPSYCH, The Lundbeck Foundation Initiative for Integrative Psychiatric Research, DK and Mental Health Center Copenhagen, Copenhagen Universtity Hospital, Copenhagen, DK), Markus M Nöthen (Institute of Human Genetics, University of Bonn, Bonn, DE and Life&Brain Center, Department of Genomics, University of Bonn, Bonn, DE), Michael C O’Donovan (MRC Centre for Neuropsychiatric Genetics and Genomics, Cardiff University, Cardiff, GB), Sara A Paciga (Human Genetics and Computational Biomedicine, Pfizer Global Research and Development, Groton, CT, US), Nancy L Pedersen (Department of Medical Epidemiology and Biostatistics, Karolinska Institutet Stockholm, SE), Brenda WJH Penninx (Department of Psychiatry, Vrije Universiteit Medical Center and GGZ inGeest, Amsterdam, NL), Roy H Perlis (Department of Psychiatry, Massachusetts General Hospital, Boston, MA, US and Psychiatry, Harvard Medical School, Boston, MA, US), David J Porteous (Medical Genetics Section, CGEM, IGMM, University of Edinburgh, Edinburgh, GB), James B Potash (Psychiatry, University of Iowa, Iowa City, IA, US), Martin Preisig (Department of Psychiatry, University Hospital of Lausanne, Prilly, Vaud, CH), Marcella Rietschel (Department of Genetic Epidemiology in Psychiatry, Central Institute of Mental Health, Medical Faculty Mannheim, Heidelberg University, Mannheim, Baden-Württemberg, DE), Catherine Schaefer (Division of Research, Kaiser Permanente Northern California, Oakland, CA, US), Thomas G Schulze (Department of Genetic Epidemiology in Psychiatry, Central Institute of Mental Health, Medical Faculty Mannheim, Heidelberg University, Mannheim, Baden-Württemberg, DE, Institute of Psychiatric Phenomics and Genomics (IPPG), Medical Center of the University of Munich, Campus Innenstadt, Munich, DE, Department of Psychiatry and Behavioral Sciences, Johns Hopkins University, Baltimore, MD, US, Department of Psychiatry and Psychotherapy, University Medical Center Göttingen, Goettingen, Niedersachsen, DE and Human Genetics Branch, NIMH Division of Intramural Research Programs, Bethesda, MD, US), Jordan W Smoller (Department of Psychiatry, Massachusetts General Hospital, Boston, MA, US, Psychiatric and Neurodevelopmental Genetics Unit (PNGU), Massachusetts General Hospital, Boston, MA, US and Stanley Center for Psychiatric Research, Broad Institute, Cambridge, MA, US), Kari Stefansson (deCODE Genetics / Amgen, Reykjavik, IS and Faculty of Medicine, University of Iceland, Reykjavik, IS), Henning Tiemeier (Epidemiology, Erasmus MC, Rotterdam, Zuid-Holland, NL, Child and Adolescent Psychiatry, Erasmus MC, Rotterdam, Zuid-Holland, NL and Psychiatry, Erasmus MC, Rotterdam, Zuid-Holland, NL), Rudolf Uher (Psychiatry, Dalhousie University, Halifax, NS, CA), Henry Völzke (Institute for Community Medicine, University Medicine Greifswald, Greifswald, Mecklenburg-Vorpommern, DE), Myrna M Weissman (Psychiatry, Columbia University College of Physicians and Surgeons, New York, NY, US and Division of Epidemiology, New York State Psychiatric Institute, New York, NY, US), Thomas Werge (iPSYCH, The Lundbeck Foundation Initiative for Integrative Psychiatric Research, DK, Institute of Biological Psychiatry, Mental Health Center Sct. Hans, Mental Health Services Capital Region of Denmark, Copenhagen, DK and Department of Clinical Medicine, University of Copenhagen, Copenhagen, DK), Cathryn M Lewis* (MRC Social Genetic and Developmental Psychiatry Centre, King’s College London, London, GB and Department of Medical & Molecular Genetics, King’s College London, London, GB), Douglas F Levinson (Psychiatry & Behavioral Sciences, Stanford University, Stanford, CA, US), Gerome Breen (MRC Social Genetic and Developmental Psychiatry Centre, King’s College London, London, GB and NIHR BRC for Mental Health, King’s College London, London, GB), Anders D Børglum (Department of Biomedicine, Aarhus University, Aarhus, DK, iSEQ, Centre for Integrative Sequencing, Aarhus University, Aarhus, DK and iPSYCH, The Lundbeck Foundation Initiative for Integrative Psychiatric Research, DK), Patrick F Sullivan (Department of Medical Epidemiology and Biostatistics, Karolinska Institutet Stockholm, SE, Genetics, University of North Carolina at Chapel Hill, Chapel Hill, NC, US and Psychiatry, University of North Carolina at Chapel Hill, Chapel Hill, NC, US). **PGC-MDD group Chairs*:** Catherine Lewis and Andrew McIntosh.

## Supporting information

### S1 Supporting Information

**S1 Fig. Genetic stress-sensitivity effect representation.** Genetic stress-sensitivity effect on MDD (*β*_SS_) is defined as the difference between the regression coefficient in MDD cases (*β*_A_) and the regression coefficient in controls (*β*_B_) from linear models regressed on EPQN, adjusted by covariates. *A1:* allele 1. *A2* allele 2.

**S2 Fig. QQ plot from stress-sensitivity meta-analysis.** QQ plot of GWIS from sample size weighted meta-analysis (λ = 0.997; s.e. = 1.05×10^-5^). All SNPs wit *p* < 2×10^-5^, *p* threshold (dot line) where some SNPs start to deviate from null distribution going outside 95% confidence intervals (grey shadow), were selected to perform DEPICT analyses to assess pathway and functional genomic analyses. 27 top variants from 12 independent loci were selected.

**S3 Fig. QQ plots of GWIS *p* values.** QQ plots of GWIS from (A) UKB (λ = 1.014; s.e.= 1.027×10^-5^), (B) GS:SFHS (λ = 0.997; s.e. = 7.989×10^-6^). The 95% confidence interval is shaded in grey.

**S4 Fig. Miami plots on UK Biobank and Generation Scotland: Scottish Family Health Study.** Miami plots showing comparison between association profile between SS and MDD main additive effects. Miami plots from (A) UKB filtering for SS *p* values (top) and MDD *p* values (bottom), (B) GS:SFHS filtering for SS *p* values (top) and MDD *p* values (bottom). Filter at *p* = 1×10^-3^. The x-axis is base-paired chromosomal position and y-axis is the significance (-log10p) of association with (up; red dots) SS effect and (down; blue dots) MDD. Dot line: genome-wide suggestive threshold (*p* = 1×10” ^5^) at the filtered effect; dashes lines: *p* value = 0.01 and 0.05 at compared effect.

**S5 Fig. Manhattan plot of the gene-based test for stress-sensitivity.** Manhattan plot showing gene-based association of stress-sensitivity. The x-axis is base-paired chromosomal position and y-axis is the significance (-log_10_ *p* value) of association with SS effect. Genome-wide significance threshold showed by red dashed line was defined at *p* = 0.05/17,931 = 2.79×10^-6^.

**S6 Fig. PRS profiling predicting MDD in UK Biobank.** MDD risk explained (R^2^ coefficient (%); top bar values) on the liability scale by each PRS in UKB; weighted by GWAS main additive and GWIS stress-sensitivity effects independently and combined. (A) Using summary statistics from GS:SFHS as discovery sample. (B) Replication fitting PRS_D_ and PRS_N_ using summary statistics from worldwide consortiums (i.e. PGC & GPC). Significance codes: *p* values *** < 0.001 < ** < 0.01 < * < 0.05; derived from likelihood ratio tests. *SS* stands for stress-sensitivity.

**S1 Table. EPQN comparison between MDD cases and healthy controls.**

**S2 Table. Top 10 SNPs from GWIS on UK Biobank.**

**S3 Table. Top 10 SNPs from GWIS on Generation Scotland: Scottish Family Health Study.**

**S4 Table. Traits with significant evidence of association with closest gene to suggestive stress-sensitive hits.** The closest genes to SNPs associated with stress-sensitivity at suggestive significance levels have prior evidence of association in dbGAP with a wide range of neuropsychiatric traits such as schizophrenia, bipolar disorder, attention deficit disorder with hyperactivity, mental competency, intuition, sleep or alcohol drinking.

**S5 Table. Top 25 hits from gene-based analysis of GWIS meta-analysis.**

**S6 Table.: Summary results from polygenic risk score (PRS) analysis using PRSice-2.**

**S7 Table. MDD stratification.**

